# Pangenome and transcriptome analyses reveal extensive diversity and breeding signatures in the oat GDSL-esterase/lipase gene family

**DOI:** 10.64898/2026.09.28.755230

**Authors:** Tuomas Hämälä, Sirja Viitala

## Abstract

Lipolytic enzymes play critical roles in plant development, growth, and stress responses. In oat (*Avena sativa*), they are of particular interest because lipid hydrolysis contributes to rancidity and reduced grain quality, yet little is known about genomic variation in oat lipolytic enzymes. Here, we investigated genomic and expression diversity of the GDSL-esterase/lipase gene family using pangenome, transcriptome, and genome-wide association analyses. Across 22 *A. sativa* and two *A. sterilis* genome assemblies, we identified 7,775 GDSL genes (~324 per assembly), revealing wide conservation of ancestral orthologs alongside extensive copy-number variation among homoeologs and paralogs. Transcriptome profiling across 20 genotypes and six tissues demonstrated preferential expression in embryo and root, with tissue-specific patterns highlighting candidates for specific functions, such as seed lipid metabolism. Associations with structural variants further uncovered candidate mutations underlying expression variation among accessions. Comparisons of cultivated and wild oat detected loci with signatures of breeding-based selection, suggesting that GDSL genes have contributed to diversification of the cultivated gene pool. Using 629 whole-genome sequenced accessions, we also identified associations between GDSL regions and agronomic traits, particularly plant height and effective tiller number. Several regions showed patterns consistent with selection, highlighting candidate loci for further investigation of agronomic variation. Together, our results reveal substantial genomic and transcriptional diversity within the oat GDSL gene family and identify candidate genes and genomic regions for improving seed quality and agronomic performance. More broadly, our study demonstrates how pangenomic resources can be used to investigate gene-family diversity and function in complex polyploid crops.

## Introduction

GDSL-esterase/lipases are a large and diverse family of lipolytic enzymes defined by a conserved GDSL (GDSxxDxG) sequence motif near their N-terminus. Unlike classical lipases, which typically contain a GxSxG motif, GDSL enzymes belong to the SGNH-hydrolase superfamily and possess a structurally flexible active site with four catalytic residues (Ser, Gly, Asn, and His) distributed across distinct consensus blocks (Akoh et al. 2004). This structural flexibility enables GDSL proteins to accommodate a broad range of substrates, and they have been implicated in diverse biological processes, including lipid metabolism, seed development, cuticle formation, and responses to biotic and abiotic stresses (Shen et al. 2022; Dong et al. 2026). GDSL genes have been studied in several plant species (Chepyshko et al. 2012; Lai et al. 2017; Cenci et al. 2022; Ding et al. 2023), but their diversity and functions in oat (*Avena sativa*) remain poorly characterised. Lipolytic enzymes are particularly important for oat production, because oat grains contain high levels of lipids, whose hydrolysis results in rancidity and reduced grain quality (Lehtinen and Laakso 2004). Although GDSL proteins are not the only lipolytic enzymes in plants, their large gene family and diverse expression patterns make them potential contributors to variation in oat lipid metabolism and grain quality. At the same time, given the diverse functions of GDSL proteins in other species, members of this gene family may also contribute to a range of developmental, physiological, and agronomic processes in oat (Shen et al. 2022). Characterising their genomic and transcriptional diversity may therefore provide insights extending beyond grain lipid metabolism and rancidity.

Understanding the contribution of GDSL genes to these processes requires characterising genetic variation across a diverse set of genomes. Traditional analyses based on a single reference genome are poorly suited to this task because they primarily capture variation relative to one genotype and can overlook genes and genomic regions that are absent from that assembly.Pangenome analyses overcome this limitation by considering multiple genomes simultaneously, enabling the efficient characterisation of genomic structural variants, gene presence/absence variation, and copy-number variation (Jayakodi et al. 2025). These approaches have provided new insights into the evolution and functional diversity of individual gene families in several plant species (Barragan and Weigel 2021; Zhao et al. 2025; Tan et al. 2026; Anam et al. 2026).

Oat presents an especially interesting system for investigating pangenomic variation. *A. sativa* is a allohexaploid species with a large and structurally complex genome, containing three related subgenomes and extensive repetitive DNA (Kamal et al. 2022). This genomic complexity has historically limited the development and use of high-quality genomic resources. However, recent advances in long-read sequencing have substantially improved our ability to investigate genomic variation in complex genomes (Jayakodi et al. 2025). The increasing availability of high-quality genome assemblies from diverse cultivars, landraces, and wild species now provides an opportunity to investigate the evolutionary history and diversification of gene families across both domesticated and wild oat (Kamal et al. 2022; Peng et al. 2022; Avni et al. 2026).

Here, we use these new resources to investigate the diversity and evolution of the oat GDSL gene family. Using 24 oat genome assemblies, we characterise GDSL gene content, pangenomic variation, and expression patterns across tissues and genotypes. We further use population genetic and genome-wide association analyses to investigate signatures of selection and associations with agronomically important traits. Together, our results provide a comprehensive overview of GDSL-esterase/lipase gene-family diversity in oat and identify candidate genes and genomic regions for further investigation of their roles in grain quality and agronomic performance.

## Methods

### Identification and pangenomic characterisation of GDSL genes

To identify GDSL genes and characterise their pangenomic variation, we used high-quality genome assemblies from 22 *A. sativa* accessions and two accessions of *A. sterilis*, the wild progenitor of cultivated oat (Table S1). One *A. sativa* assembly (Marvellous) was previously published by Li et al. (2025), whereas the remaining assemblies were generated as part of the PanOat project (Avni et al. 2026). The PanOat assemblies underwent a consolidation analysis in which gene models were cross-mapped across genomes to identify and correct missing or misannotated genes, thereby improving the accuracy of gene-level variant detection. Although the Marvellous genome was not part of the consolidation analysis, it showed similar numbers and levels of variation in GDSL genes to the other assemblies (Fig. 1), supporting its inclusion in the pangenome dataset.

**Figure 1.**
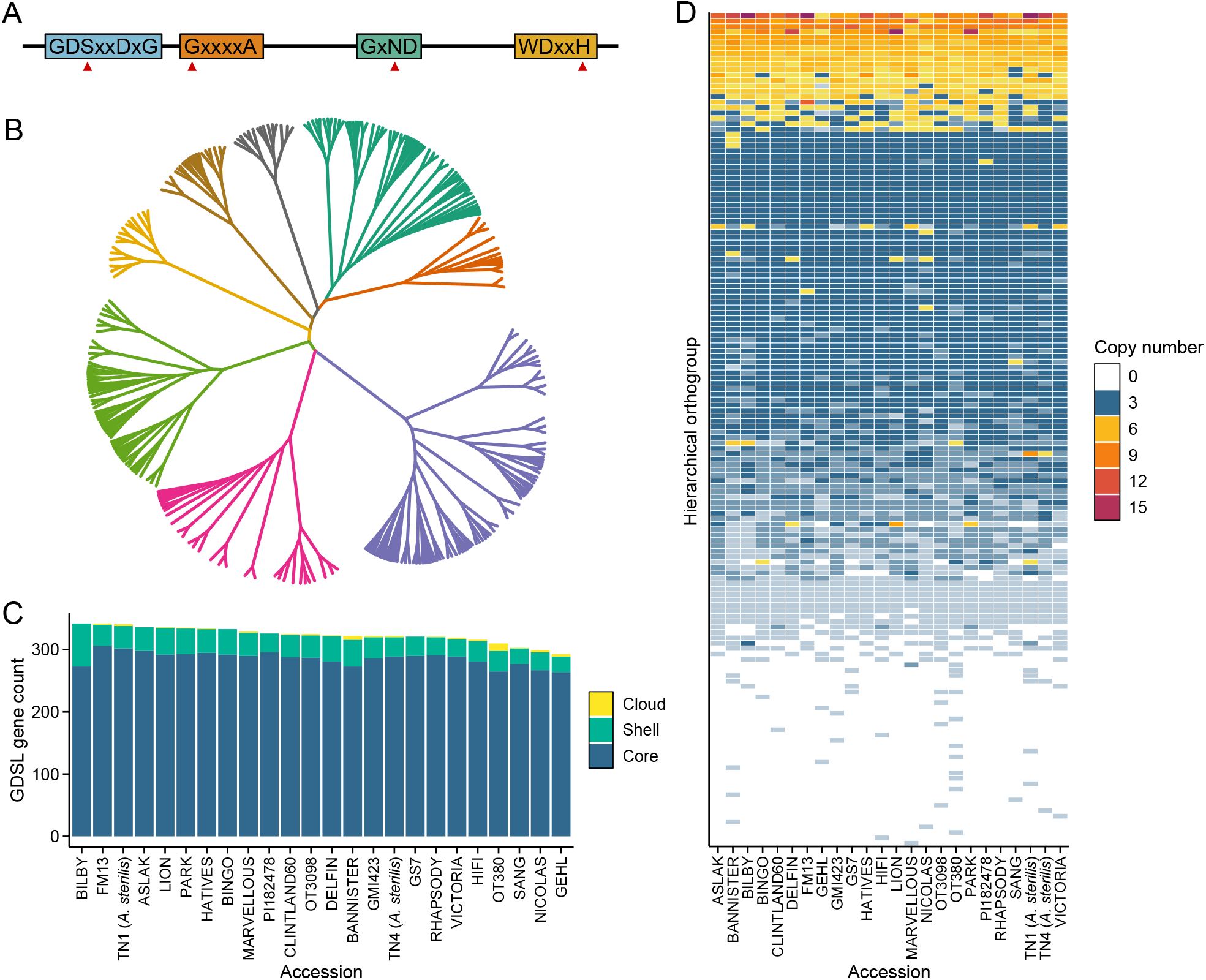
Pangenomic variation in GDSL genes. **A**: Basic structure of GDSL genes. The figure shows four conserved amino acid motifs, each containing a catalytic residue (red arrows). **B**: Maximum-likelihood tree of GDSL genes identified in the TN1 assembly. Colours indicate eight major clades. **C**: Number of GDSL genes identified in each assembly and homoeolog-aware pangenome composition. Genes were classified as core (100% presence), shell (15–<100% presence), or cloud (<15% presence) based on HOG sharing among assemblies. **D**: Copy-number variation among GDSL genes. Colours indicate the number of orthologous copies identified for each HOG–accession combination.

We chose the longest transcript for each gene with high annotation confidence and identified putative GDSL genes using the GDSL Hidden Markov Model (HMM) profile from Pfam (PF00657) and HMMER 3.0 (hmmer.org). Candidate genes were required to have an *e*-value < 1 × 10^−5^ and contain two conserved amino acid motifs, GDSxxDxG and DxxH (where x denotes any amino acid), which encompass the catalytic triad (S-D-H) characteristic of active GDSL enzymes (Akoh et al. 2004). Additional conserved motifs were identified using MEME (Bailey et al. 2009), and phylogenetic relationships among GDSL genes inferred using IQ-TREE (Wong et al. 2026).

To characterise presence/absence and copy-number variation among GDSL genes, we inferred orthologous relationships using OrthoFinder (Emms et al. 2026). Following Avni et al. (2026), genes were separated by subgenome, OrthoFinder ran with default settings, and the hierarchical orthogroups (HOGs) classified according to their frequency across the pangenome as core (100% presence), shell (15–<100% presence), or cloud (<15% presence). For each accession, we also calculated the number of genes assigned to each HOG to assess copy-number variation.

For analyses requiring genes to be matched between assemblies, the *A. sterilis* accession TN1 was used as the reference. Coding sequences were aligned using BLAST, and the best match for each gene was retained when it met the following criteria: *e*-value < 1 × 10^-10^, assignment to corresponding subgenomes, ≥90% sequence identity, and ≥80% alignment coverage.

### Transcriptome analyses

We used processed transcriptome data from Avni et al. (2026) to characterise expression patterns across GDSL genes. The dataset comprised 20 *A. sativa* accessions and six tissues (root, embryo, panicle, caryopsis, leaf, and internode), with data aggregated across nine biological replicates for each genotype–tissue combination. Gene expression levels were quantified by aligning RNA-sequencing reads from each accession to its corresponding genome assembly and normalised as transcripts per million (TPM). See Avni et al. (2026) for further details on data generation and processing.

We first used redundancy analysis (RDA) to investigate multivariate expression patterns among the GDSL genes. TPM values were log_10_-transformed, and partial RDA models fitted using the R package vegan (Oksanen et al. 2026) to partition expression variation attributable to genotype and tissue type.

Next, to assess whether genomic structural variants (SVs) contributed to between-genotype variation in GDSL expression, we identified large (>50 bp) insertions and deletions in regions surrounding the genes. For each of the 20 *A. sativa* genomes, we extracted 1-Mb blocks centred on GDSL genes using BEDtools (Quinlan and Hall 2010), collapsed overlaps, and aligned the resulting sequences against the TN1 genome using minimap2 (Li 2018). Insertions and deletions were identified and genotyped across genomes using SVGAP (Hu et al. 2025). We focused on SVs located within 10 kb of GDSL genes and searched for associations with gene expression by comparing the fit of two linear models:

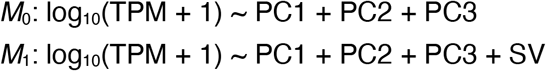

Three genomic principal components (PCs) were included as covariates to account for population structure (the number was defined using a scree plot). Differences in model fit were assessed using likelihood-ratio tests (LRTs) and *P*-values corrected for multiple testing by transforming them to false-discovery rate (FDR)-based *Q*-values (Storey and Tibshirani 2003). We then estimated the proportion of expression variation explained by each SV by calculating a pseudo-*R*^2^ as 1 − *D*(*M*_1_) / *D*(*M*_0_), where *D*(*M*_i_) denotes the residual deviance of the model. This metric thus represents the relative improvement in model fit after accounting for the SV genotype.

Tissue-specific expression patterns were then assessed both within individual genotypes and across all accessions. For analyses including all accessions, log_10_-transformed TPM values were averaged across genotypes. Tissue specificity was quantified using the tau (τ) index (Kryuchkova-Mostacci and Robinson-Rechavi 2017), with τ ≥0.8 used as the threshold for tissue-specific expression.

We last analysed subgenome dominance in complete homoeologous triads, defined using HOGs, following the approach described in Ramírez-González et al. (2018). Briefly, expression levels within each triad were compared with the idealised expression patterns expected under seven subgenome-dominance scenarios (Fig. S1). Euclidean distances between the observed and expected expression patterns were calculated, and the scenario with the shortest distance was assigned to each triad.

### Selection analyses

To identify signatures of selection associated with domestication and breeding in GDSL genes, we compared sequence variation in *A. sativa* with that of its wild progenitor, *A. sterilis*. We first estimated the ratio of non-synonymous to synonymous nucleotide divergence (*d*_N_/*d*_S_) between the two *A. sterilis* accessions and each *A. sativa* accession. Coding sequences were aligned at the codon level using PRANK (Löytynoja and Goldman 2008), and *d*_N_ and *d*_S_ calculated using the R package seqinr (Charif and Lobry 2007). Values were summed across all pairwise comparisons, and global *d*_N_/*d*_S_ ratio calculated as the ratio of the summed *d*_N_ and *d*_S_ values. 95% confidence intervals (CIs) were estimated by randomly resampling the pairwise comparisons 1,000 times.

Because *d*_N_/*d*_S_ has limited sensitivity to detect recent positive selection, we additionally searched for genes harbouring high-frequency derived alleles using Fay and Wu’s *H* (Fay and Wu 2000). Ancestral alleles were defined as nucleotides shared by both *A. sterilis* accessions. Unfolded site-frequency spectra (SFS) were then estimated for each gene, and normalised Fay and Wu’s *H* calculated from the resulting SFS. 95% CIs were estimated from 1,000 parametric bootstrap SFS generated using the R function rmultinom.

To further characterise nucleotide variation in the GDSL genes, we estimated pairwise nucleotide diversity (π) and Tajima’s *D* for the coding sequences. The functional consequences of amino acid-changing mutations with a derived allele frequency ≥0.9 were predicted using PPVED (Gou et al. 2022).

Finally, to visualise evidence of a selective sweep surrounding the candidate gene 3AG01900850, we used MUMmer (Marçais et al. 2018) to identify SNPs within a 1 Mb region centred on the gene. Chromosome 3A from each assembly was aligned against TN1 using nucmer, and SNPs and aligned regions were identified using show-snps and show-coords, respectively. SNPs were filtered to retain only sites uniquely aligned across all comparisons and derived allele frequencies estimated in 20 kb windows across the region.

### Genome-wide association analyses

We assessed potential phenotypic effects of GDSL variants using genome-wide association (GWA) analyses. SNP and phenotypic data for 629 *A. sativa* accessions were obtained from OatOmics (Li et al. 2025). The SNP calls were based on whole-genome resequencing data aligned to the Marvellous genome assembly. Phenotypic data were available for eight agronomic traits, measured in a field experiment in China: plant height, stem diameter, tiller number, effective tiller number, flag leaf length, flag leaf width, fresh weight, and dry weight.

GWA analyses were performed using univariate linear mixed models implemented in GEMMA (Zhou and Stephens 2012). Only SNPs located within 10 kb of GDSL genes, with a minor allele frequency (MAF) >0.05 and <10% missing data, were included. To account for population structure and cryptic relatedness, four principal components (PCs) were included as fixed-effect covariates and a genetic relationship matrix (GRM) included as a random effect. PCs and the GRM were estimated from linkage-pruned SNP data (*r*^2^ ≤ 0.1 within a 100-SNP window) using PLINK (Chang et al. 2015) and GEMMA, respectively. The number of PCs included as covariates was determined using a scree plot. For each phenotype, *P*-values were corrected for multiple testing by converting them to FDR-based *Q*-values (Storey and Tibshirani 2003).

## Data availability

All data used in this study were previously published and are publicly available (Li et al. 2025; Avni et al. 2026). Custom scripts used for the analyses are available at: https://github.com/thamala/gdsl.

## Results

### GDSL genes exhibit presence/absence and copy-number variation in the oat pangenome

Using a GDSL HMM profile together with the presence of two diagnostic amino acid motifs, we identified 7,775 putative GDSL genes across the 24 oat assemblies, corresponding to an average of 324 genes per assembly. Searches for additional conserved amino acid motifs confirmed the presence of four catalytic residues characteristic of GDSL genes (Fig. 1A). To examine the phylogenetic relationships among the identified genes, we aligned all GDSL protein sequences from the *A. sterilis* assembly TN1 and constructed a maximum-likelihood phylogeny. The tree revealed distinct sequence-based clusters, consistent with substantial diversification among GDSL genes, and we divided the genes into eight clades based on the major branching patterns (Fig.1B).

We next used OrthoFinder (Emms et al. 2026) to identify sets of genes across the pangenome that originated from common ancestral genes. The resulting hierarchical orthogroups (HOGs) comprised both single-copy orthologs, representing genes without duplication events within the studied lineages, and in-paralogs, resulting from duplications that occurred after lineage divergence. Note that here we analysed each subgenome separately, meaning that single-copy orthologs can still have up to three copies (homoeologs) in each accession. OrthoFinder identified 151 HOGs that were largely shared by all accessions and therefore classified as core, whereas only 12% of GDSL genes belonged to HOGs in the dispensable group (i.e., shell and cloud; Fig. 1C). Despite this high level of HOG sharing, we observed extensive copy-number variation, with 66% of HOG–accession combinations deviating from the expected three copies (Fig. 1D). The extent of copy-number variation was further reflected in the markedly smaller core gene fraction (~40%) obtained when homoeologs were not distinguished in the OrthoFinder analysis (Fig. S2).

### Tissue-specific expression patterns of GDSL genes

Analysis of GDSL gene expression across 20 *A. sativa* genotypes and six tissues revealed substantial variation among samples. Redundancy analysis (RDA) showed that samples clustered primarily by tissue type, whereas differences among genotypes were considerably less pronounced (Fig. 2A). Indeed, partial RDA models indicated that genotype alone explained only 4% of total expression variation, compared to 84% explained by tissue type. Nevertheless, several genes had disproportionately large loadings in the RDA models (black circles in Fig. 2A), indicating that the expression of individual GDSL genes can vary substantially between genotypes.

**Figure 2.**
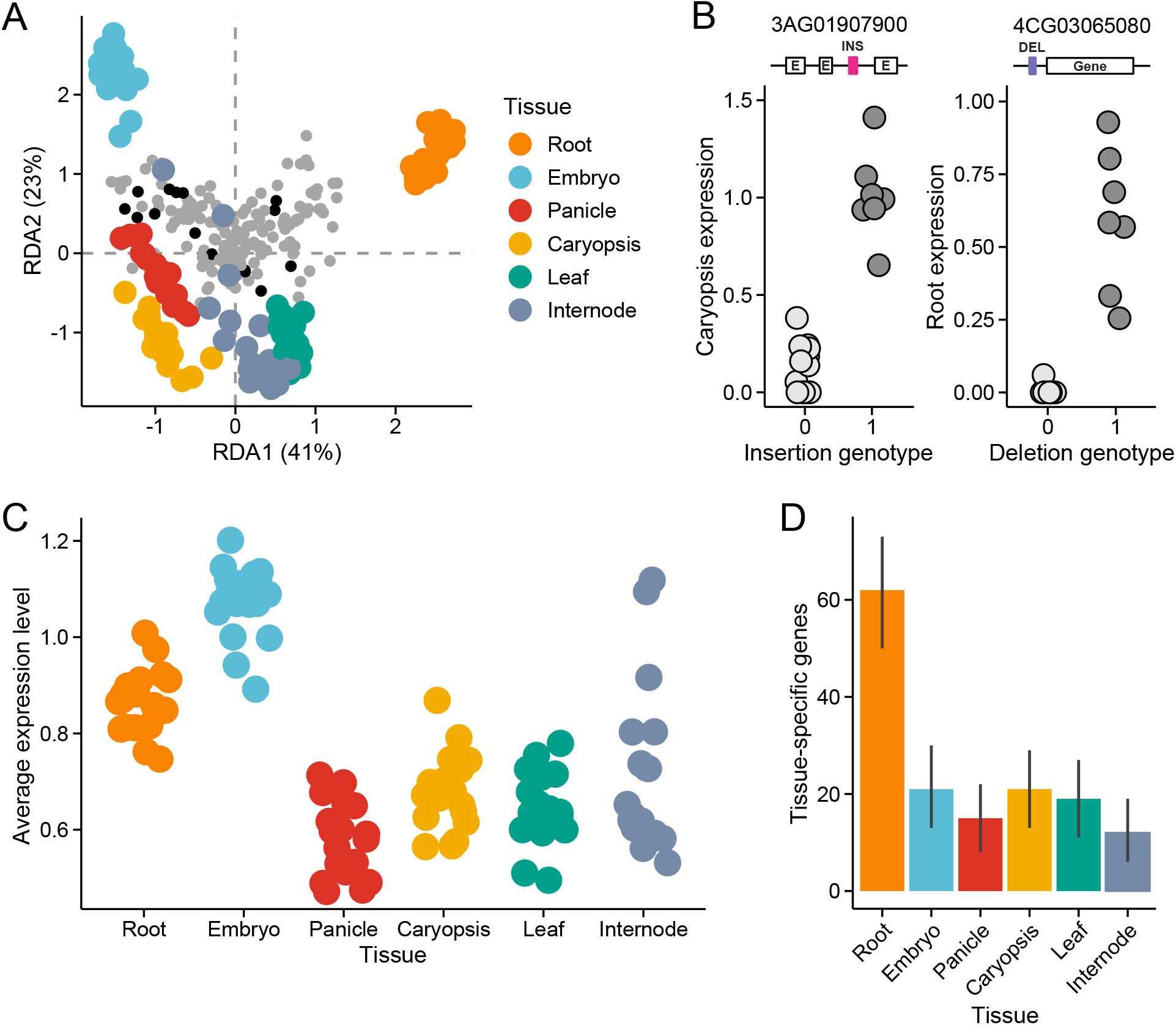
Transcriptomic variation in GDSL genes. **A**: RDA plot showing expression variation among samples. Coloured circles indicate tissue types, while gene loadings are shown in the background. Black circles indicate genes that were outliers in a genotype-based model and therefore explained a relatively large proportion of between-genotype variation. The proportion of variance explained by each RDA axis is shown in parentheses. **B**: Allelic effects of SVs found in the regulatory regions of two genes, 3AG01907900 and 6CG04833700. Schematics show the location of the insertion (INS) and deletion (DEL) in relation to the gene or exons (marked with letter ‘E’) in the TN1 assembly. Vertical axes show log_10_-transformed TPM values in caryopsis and root. **C**: Expression levels of GDSL genes across accessions and tissues. Circles indicate mean log_10_-transformed TPM values for each accession. **D**: Number of tissue-specific genes identified in each tissue. Error bars indicate 95% CIs. Expression data were aggregated across genotypes (accession-specific results are shown in Fig. S3).

To identify genomic variants that may contribute to this variation, we examined structural variants (SVs; >50 bp) located in *cis-*regulatory regions around GDSL genes (introns, 10 kb up- and downstream). As SVs typically have a large influence on local genomic structure, and often contain regulatory elements associated with transposable elements, they tend to have an outsized impact on the expression of nearby genes (Alonge et al. 2020; Hämälä et al. 2021) Out of 157 SV–gene pairs with sufficient variation, we identified 14 significant expression associations at *Q* < 0.05.

Notably, a 222-bp insertion in the intron of 3AG01907900 explained up to 86% of expression variation in caryopsis, while a 179-bp deletion 370 bp upstream of 6CG04833700 explained up to83% of expression variation in root (Fig. 2B). These findings indicate that segregating SVs can contribute substantially to GDSL expression variation among cultivated oat accessions.

Given the strong effect of tissue type on GDSL gene expression, we next analysed each tissue separately. The highest GDSL expression levels were observed in embryo and root, whereas panicle showed the lowest expression (Fig. 2C). Because mean expression levels provide only a general indication of tissue affinity, we additionally identified genes with preferential expression in individual tissues. Roots contained the largest number of tissue-specific GDSL genes, whereas the other tissues showed broadly similar numbers (Fig. 2D). Despite these overall similarities, individual tissue-preferential genes may represent candidates underlying tissue-specific biological functions, such as seed lipid metabolism and rancidity (Table S2).

The number of tissue-specific genes also varied among genotypes, ranging from 99 to 148 per accession. However, the relative distribution among tissues was broadly consistent across genotypes (Fig. S3). Finally, analysis of homoeolog expression revealed broadly balanced expression among most complete triads (Fig. S1), suggesting that the majority of retained GDSL homoeologs remain transcriptionally active in oat.

### Variation in GDSL genes has been shaped by domestication and breeding

We next examined nucleotide variation in GDSL genes to determine whether individual genes show signatures consistent with selection during oat domestication and breeding. Comparisons between cultivated *A. sativa* and two *A. sterilis* accessions (representing the wild progenitor of *A. sativa*) showed that the protein-coding sequences have predominantly evolved under purifying selection, as the ratio of nonsynonymous to synonymous divergence (*d*_N_/*d*_S_) exceeded one for only two genes (Fig. 3A). Because *d*_N_/*d*_S_ is relatively insensitive to recent positive selection and is generally not informative about selection acting outside nonsynonymous sites, we additionally used Fay and Wu’s *H* (Fay and Wu 2000) to identify genes with an excess of high-frequency derived alleles in *A. sativa*. Several genes showed negative Fay and Wu’s *H* values, indicative of recent selection on the cultivated oat lineage (Fig. 3B). Many of these genes also showed reduced nucleotide diversity and negative Tajima’s *D* values (Fig. S4), providing additional evidence for recent selective processes.

**Figure 3.**
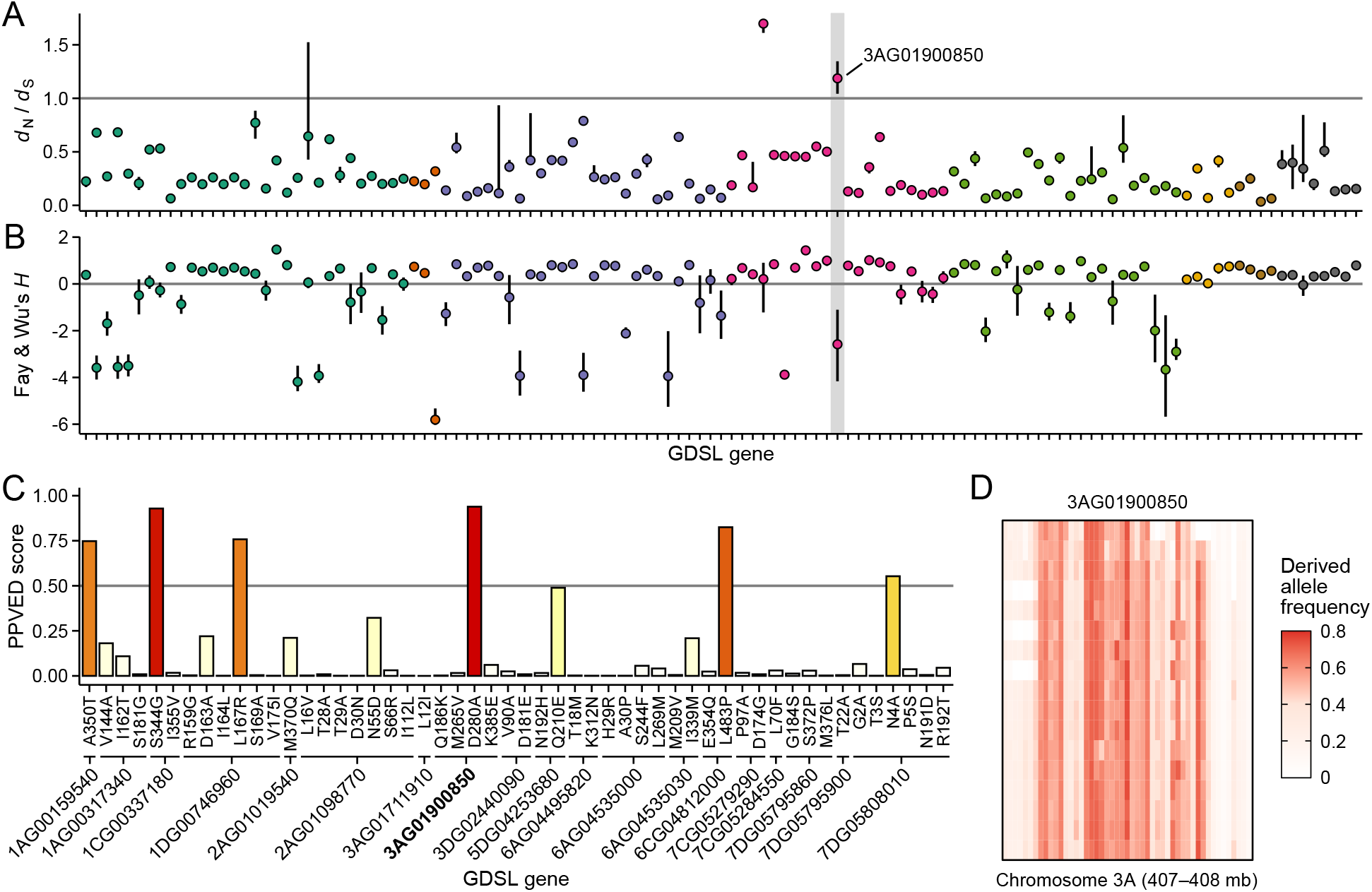
Signals of breeding-based selection in GDSL genes. **A**: Ratio of nonsynonymous to synonymous nucleotide divergence between *A. sativa* and *A. sterilis* (*d*_N_/*d*_S_). **B**: Fay and Wu’s *H* estimated from coding sequences. Negative values indicate excess of derived alleles in *A. sativa*. In panels A and B, colours correspond to the clades shown in Fig. 1B, and error bars indicate 95% CIs. **C**: Predicted functional effects of amino acid substitutions with a derived allele frequency ≥0.9. Substitutions with PPVED scores >0.5 were considered likely to have functional effects. Gene names correspond to the TN1 genome assembly. **D**: SNP variation within a 1-Mb region surrounding 3AG01900850. For each accession, derived allele frequencies were estimated in 20-kb sliding windows.

One gene, 3AG01900850, was particularly notable because it combined a relatively high *d*_N_/*d*_S_ ratio with a strongly negative Fay and Wu’s *H*, suggesting that positive selection may have contributed to the evolution of its coding sequence in cultivated oat. To investigate whether amino acid substitutions in this and other GDSL genes are likely to affect protein function, we used PPVED (Gou et al. 2022) to predict the functional consequences of amino acid substitutions based on evolutionary conservation across plant species. One fixed substitution in 3AG01900850 had a high PPVED score, indicating that the mutation may affect protein function (Fig. 3C). PPVED identified five additional highly diverged substitutions in other GDSL genes with predicted functional effects (score > 0.5; Fig. 3C), suggesting that novel GDSL gene products may have been selected for in domesticated oat.

The patterns observed at 3AG01900850 were also consistent with selection acting relatively early during the domestication of oat, as all *A. sativa* accessions carried the derived allele at multiple positions across the gene. To further investigate whether the surrounding genomic region showed evidence of a selective sweep, we examined SNP variation within 1 Mb of 3AG01900850 and estimated derived allele frequencies across cultivated oat accessions. The resulting SNP matrix revealed blocks of high-frequency derived alleles centred on the gene (Fig. 3D), consistent with a hard selective sweep. However, given the extensive linkage disequilibrium across this region, these data do not allow us to determine whether the amino acid substitution itself was the target of selection or whether the sweep was driven by a linked variant.

### GDSL genes are associated with phenotypic variation in agronomic traits

To investigate potential phenotypic effects of GDSL variants, we conducted genome-wide association (GWA) analyses using a globally distributed panel of 629 resequenced *A. sativa* accessions (Fig. 4A), representing a diverse collection of cultivars and landraces (Fig. 4B). Of the eight agronomically important traits analysed, significant associations were detected for two traits at *Q* < 0.05: plant height and effective tiller number (Fig. 4C). The associated signals comprised five QTL regions that either overlapped or were located within 10 kb of GDSL genes (Fig. 4D).

**Figure 4.**
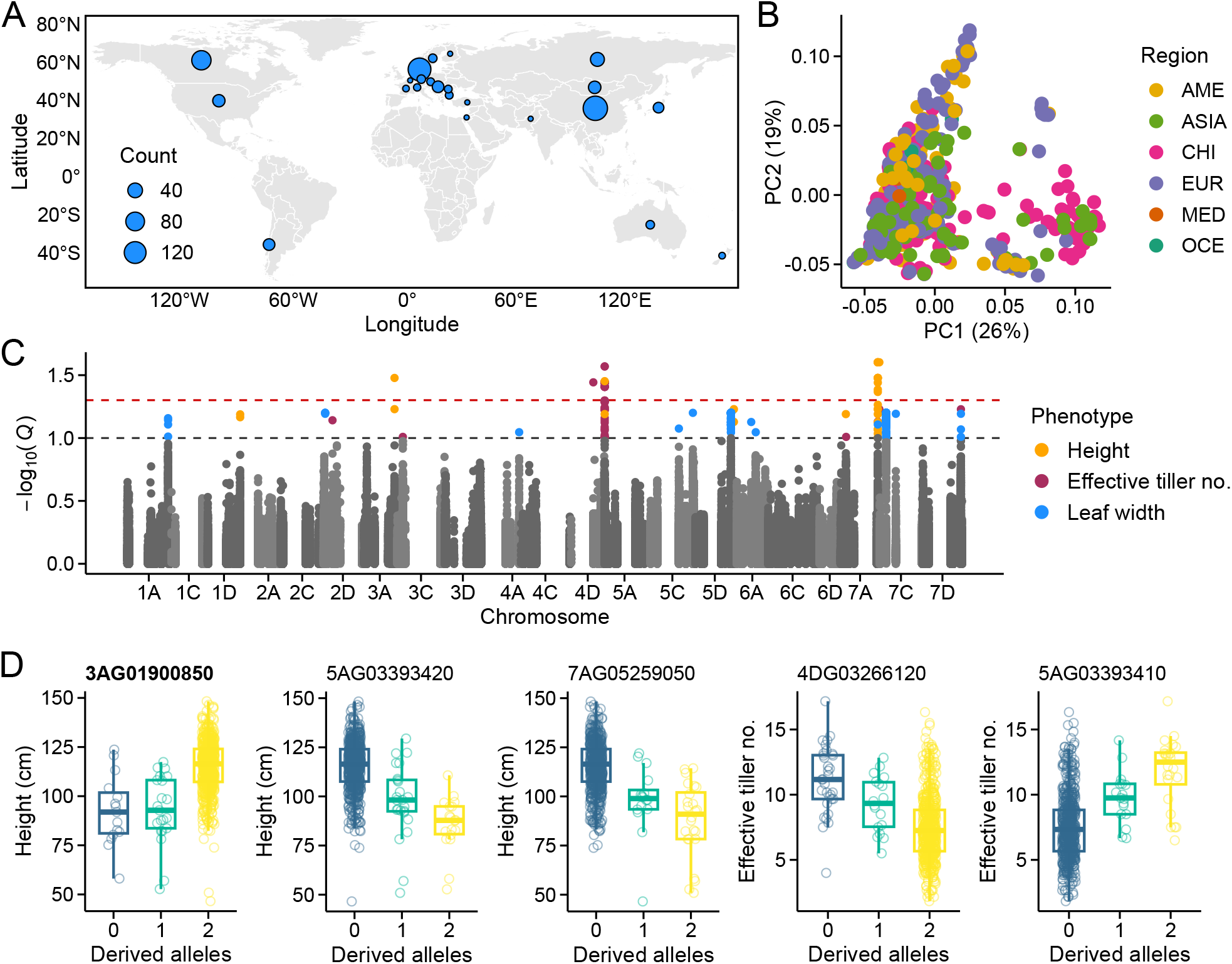
Phenotypic associations with GDSL genes. **A**: Geographic distribution of 629 *A. sativa* accessions used in the association analyses. **B**: First two principal components (PCs) from a principal component analysis. The proportion of variation explained by each PC is shown in parentheses. **C**: *Q*-values from genome-wide association (GWA) analyses of eight traits using SNPs located within 10 kb of GDSL genes. Coloured circles indicate SNPs associated with the respective traits at *Q* < 0.1. **D**: Allelic effects of five QTL detected at *Q* < 0.05 (red horizontal line in panel C). Phenotypes are shown for accessions carrying different genotypes of the lead SNP at each QTL.

One plant-height QTL was located upstream of 3AG01900850, the gene that showed evidence of breeding-based selection in our analyses (Fig. 3). Derived alleles were found at high frequency within this QTL, consistent with selection acting on the cultivated oat lineage (Fig. 4D). However, accessions carrying the *A. sativa*-type alleles were taller than those carrying the *A. sterilis*-type alleles. This contrasts with the general direction of selection for reduced plant height during cereal domestication and breeding (Miao et al. 2024), suggesting that the association may reflect pleiotropic or linked effects rather than direct selection for increased plant height at this locus.

A second QTL, located adjacent to 4DG03266120, also contained high-frequency derived alleles, providing a potential signal of selection. The coding region of 4DG03266120 was invariant across our pangenome dataset, precluding the detection of coding-sequence selection signatures at this locus. However, analysis of the short-read OatOmics dataset (Li et al. 2025) revealed substantial allele-frequency differentiation between *A. sativa* and *A. sterilis* upstream of the gene (Fig. S5), consistent with selection in the cultivated lineage. However, given the large size of the associated haplotype block, the actual target of selection may have been another gene linked to 4DG03266120.

For the remaining three QTL, derived alleles were present at low frequency and were associated either with reduced plant height or increased effective tiller number (Fig. 4D), both of which are generally favourable agronomic characteristics targeted by cereal breeding (Miao et al. 2024; Mmbando 2025). Together, these associations indicate that variation surrounding several GDSL genes coincides with phenotypic variation in traits of agronomic importance and, in some cases, with genomic patterns consistent with selection during oat domestication and breeding.

## Discussion

### Extensive pangenomic diversity of the oat GDSL gene family

GDSL-esterase/lipases constitute large gene families in plants, with previous studies identifying 105, 114, and 240 GDSL genes in *Arabidopsis thaliana*, rice, and rapeseed (Chepyshko et al. 2012; Lai et al. 2017; Ding et al. 2023). Here, we identified more than 7,000 GDSL genes across 24 oat genome assemblies, corresponding to more than 300 genes per assembly despite applying a relatively stringent identification protocol. The family size is consistent with the large, hexaploid oat genome, where ancestral loci are often represented by multiple homoeologous copies, while also providing ample opportunities for gene duplication and retention.

Previous studies of plant GDSL families have largely relied on single reference genomes, limiting their ability to capture variation among individuals. By analysing multiple oat assemblies, we identified substantial variation in GDSL gene content that would not be apparent from a single reference. Most GDSL genes belonged to HOGs broadly shared across the pangenome, yet individual groups frequently deviated from the expected three copies per accession. Thus, GDSL genes are highly conserved at the ortholog level while showing extensive copy-number variation among accessions. This pattern indicates that conservation of ancestral orthologous groups coexists with substantial variation in homoeolog and paralog copy number, highlighting copy-number turnover as an important component of GDSL diversity in oat.

Our results further demonstrate the value of accounting for homoeology in polyploid pangenome analyses. When homoeologs were not distinguished, only approximately 40% of genes belonged to the core group, compared with almost 90% in the homoeolog-aware analysis. Together, these results show that pangenomic approaches can capture important levels of gene-family variation that are difficult to resolve using single-reference analyses.

### Tissue-specific expression suggests diverse functions of GDSL genes

The broad distribution of GDSL expression across tissues is consistent with the diverse functions attributed to this family in other plant species. Although tissue type explained most of the overall expression variation, individual genes showed pronounced between-genotype variation. Our analysis identified several SVs associated with this expression variation, providing candidate regulatory mutations that may contribute to differences in GDSL expression among accessions. These results highlight the value of combining pangenomic SV analyses with transcriptome data, as expression differences may be associated with genomic variants that are difficult or impossible to identify using short-read based methods (Jayakodi et al. 2025)

GDSL genes exhibited particularly high expression in embryo, consistent with reported roles in seed development, maturation, and germination (Shen et al. 2022). This pattern is also relevant to oat grain quality because lipid hydrolysis and subsequent rancidity are important issues during grain storage. Embryo- and caryopsis-preferential genes therefore represent candidates for investigating the genetic basis of lipid metabolism and rancidity, while naturally occurring variation in these genes could potentially provide useful material for breeding programmes targeting seed quality.

Roots also showed relatively high GDSL expression and contained the largest number of tissue-specific genes. GDSL proteins have been implicated in cell-wall modification and responses to pathogens and environmental stress (Zhang et al. 2017; Dong et al. 2026), processes that are particularly relevant to roots because of their interactions with soil microorganisms and exposure to variable water and nutrient availability. Although our data do not establish specific functions, the strong root expression and tissue specificity provide candidates for investigating the roles of GDSL proteins in root development and plant–soil interactions.

### Selection has shaped variation in a subset of GDSL genes

Comparison of cultivated *A. sativa* with wild *A. sterilis* identified several GDSL genes with signatures of selection in the cultivated lineage. Most genes showed patterns consistent with purifying selection, whereas a smaller subset displayed signals of positive selection. Because our analysis used a globally diverse set of cultivated accessions, these signals likely represent selection broadly shared across cultivated oat, while additional analyses of geographically structured populations could reveal further loci affected by local adaptation or breeding.

The strongest candidate, 3AG01900850, combined elevated *d*_N_/*d*_S_ and strongly negative Fay & Wu’s *H*. It also contained a fixed nonsynonymous substitution with a high predicted functional effect, while the surrounding region showed a pattern consistent with a selective sweep. Together, these observations make 3AG01900850 a strong candidate for selection affecting GDSL function. However, extensive linkage disequilibrium prevents us from determining whether the nonsynonymous substitution itself was the target of selection, highlighting the need for functional validation.

Interestingly, the region surrounding 3AG01900850 was also associated with plant height. However, cultivated-oat alleles were associated with taller plants, contrary to the general selection for reduced stature during cereal domestication and breeding (Miao et al. 2024). The association may therefore reflect pleiotropy or linkage to the actual target of selection, illustrating the difficulty of assigning phenotypic targets to selection signatures based solely on associations with a limited set of traits.

### GDSL-associated regions contribute to agronomic variation

Given the diverse functions of GDSL proteins, we tested whether variation in or near GDSL genes was associated with eight agronomically important traits. Significant associations were detected for plant height and effective tiller number, with five QTL overlapping or located close to GDSL genes. These traits are plausible candidates for GDSL-associated variation because plant height involves cell expansion and cell-wall properties, while tiller production reflects shoot development and reproductive potential (Zhang et al. 2017; Mmbando 2025).

For most QTL, minor alleles were associated with agronomically favourable phenotypes, either reduced plant height or increased effective tiller number. Although the causal variants remain unknown, these associations indicate that naturally occurring variation at GDSL genes may provide useful genetic diversity for oat breeding.

The relationship between GDSL genes and the loci showing selection signals is less straightforward. Both 3AG01900850 and the region surrounding 4DG03266120 showed evidence consistent with selection and were associated with phenotypic variation, but neither result demonstrates that the measured traits were the targets of selection. Pleiotropy or linkage to another selected locus are both possible explanations. Indeed, the traits most directly targeted during domestication and breeding may not have been included in our GWA analyses, Given the potential roles of GDSL proteins in lipid metabolism, seed development, and defence, yield and grain-quality traits could provide stronger phenotypic links to some of the selection signals identified here.

## Conclusions

Overall, our analyses revealed a large and highly variable GDSL-esterase/lipase gene family in oat. Strong conservation of orthologous groups coexists with extensive variation in gene copy number, while SVs in regulatory regions provide an additional source of genomic variation associated with GDSL expression. Tissue-specific expression points to potentially important roles in seed and root biology, while comparisons between cultivated and wild oat identified several loci with patterns consistent with selection. Associations with plant height and effective tiller number further highlighted GDSL-associated regions as potential sources of useful agronomic variation. More broadly, this study demonstrates how modern genomic resources, including pangenomes, structural variant datasets, transcriptomes, and large-scale resequencing panels, can be integrated to investigate gene-family diversity, regulation, evolution, and agronomic variation in oat, a crop that has historically lacked the genomic resources available for many other major cereals. Together, these findings provide a foundation for investigating the contributions of GDSL genes to grain quality, adaptation, and oat improvement.

## Acknowledgements

The authors wish to acknowledge CSC – IT Center for Science, Finland, for computational resources. This work was supported by Business Finland (FABulOATS, Functionality and quality-enhanced oats and faba beans for food industry).

