## Supplementary information for "Pangenome and transcriptome analyses reveal extensive diversity and breeding signatures in the oat GDSL-esterase/lipase gene family"

**Table S1.** Accession used in the pangenome analyses.

| **Accession** | **Origin** | **Type** |
|---|---|---|
| ASLAK | Finland | Cultivar |
| BANNISTER | Australia | Cultivar |
| BILBY | Australia | Cultivar |
| BINGO | Poland | Cultivar |
| CLINTLAND60 | USA | Cultivar |
| DELFIN | Germany | Cultivar |
| FM13 | Spain | Landrace |
| GEHL | Canada | Cultivar |
| GMI423 | USA | Cultivar |
| GS7 | USA | Cultivar |
| HATIVES | Switzerland | Landrace |
| HIFI | USA | Cultivar |
| LION | Germany | Cultivar |
| MARVELLOUS | Denmark | Cultivar |
| NICOLAS | Canada | Cultivar |
| OT3098 | Canada | Breeding line |
| OT380 | Canada | Breeding line |
| PARK | USA | Cultivar |
| PI182478 | Turkey | Landrace |
| RHAPSODY | UK | Cultivar |
| SANG | Sweden | Cultivar |
| TN1 | Turkey | Wild oat (*A. sterilis*) |
| TN4 | Turkey | Wild oat (*A. sterilis*) |
| VICTORIA | Argentina | Landrace |

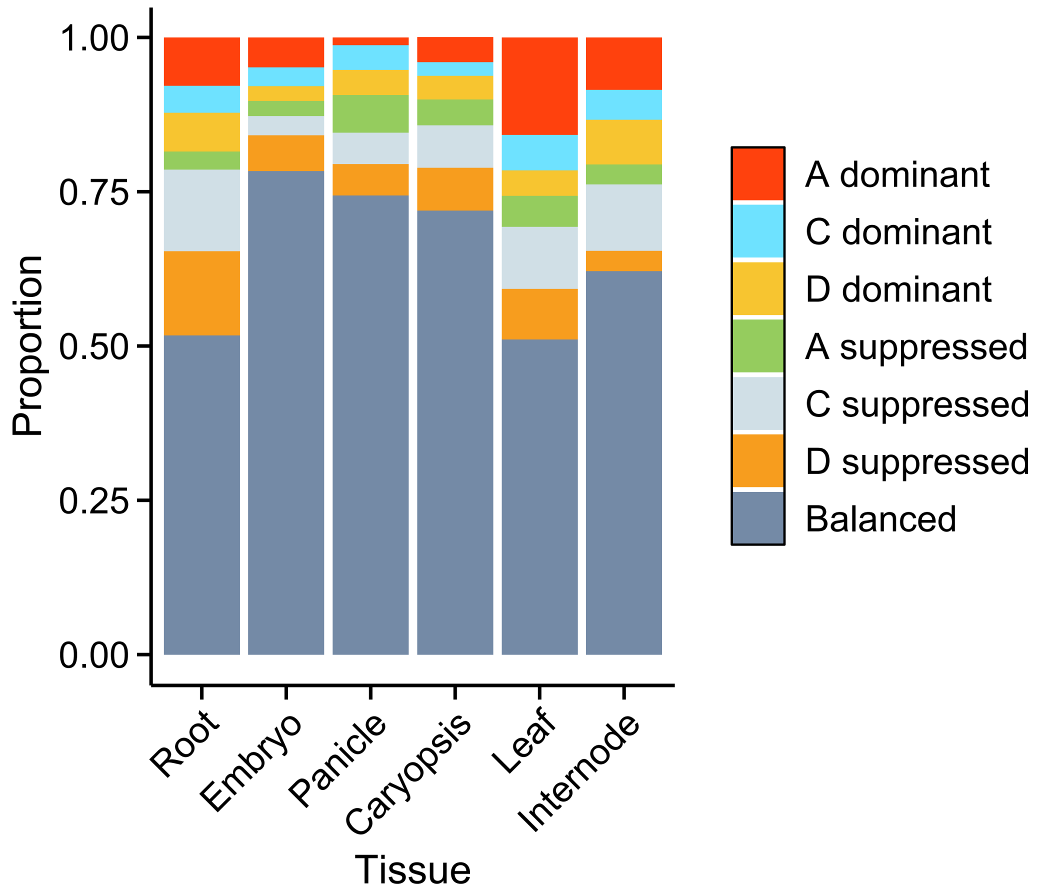


**Figure S1.** Subgenome dominance in complete homoeologous triads.

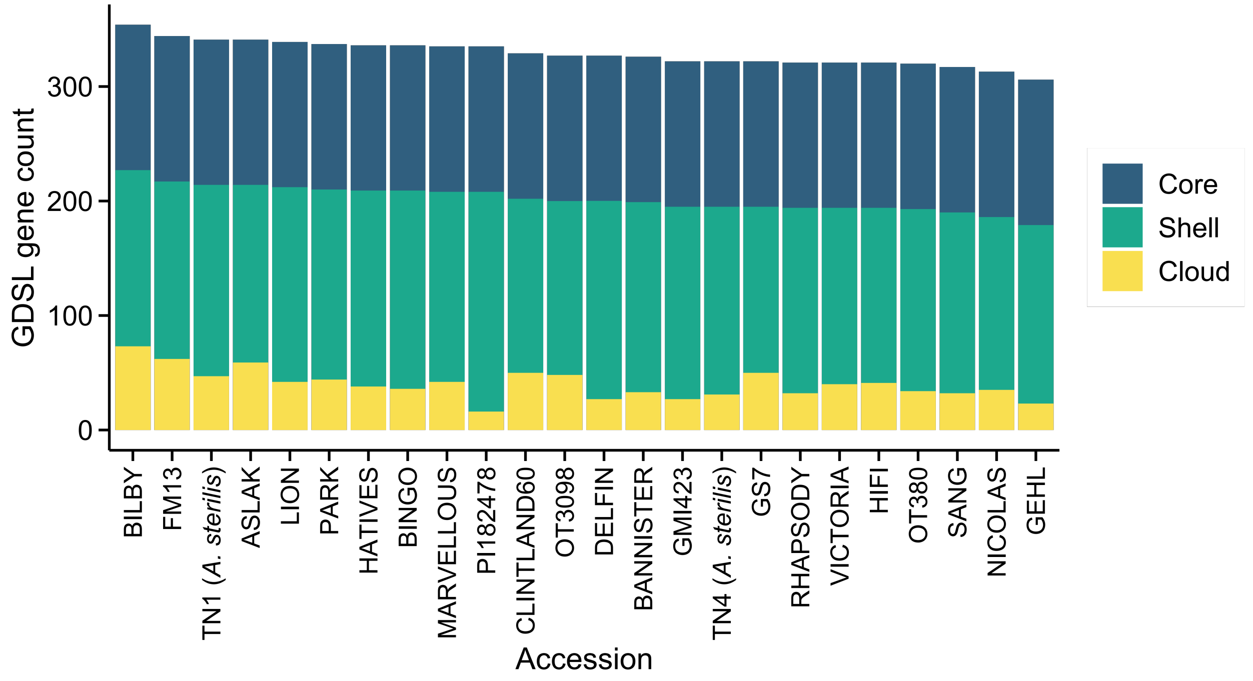


**Figure S2.** Pangenome composition when homoeologs were not considered. Genes were classified as core (100% presence), shell (15–<100% presence), or cloud (<15% presence) based on HOG sharing among assemblies.

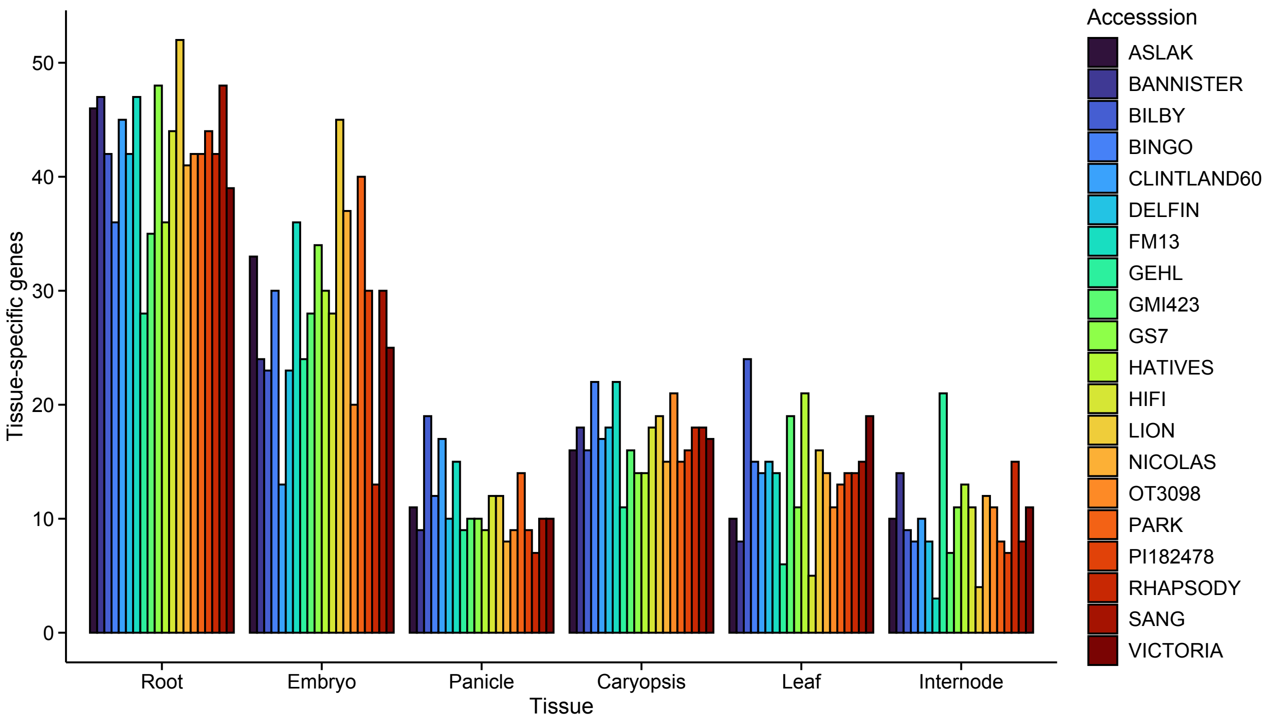


**Figure S3.** Number of tissue-specific genes identified in each tissue and genotype.

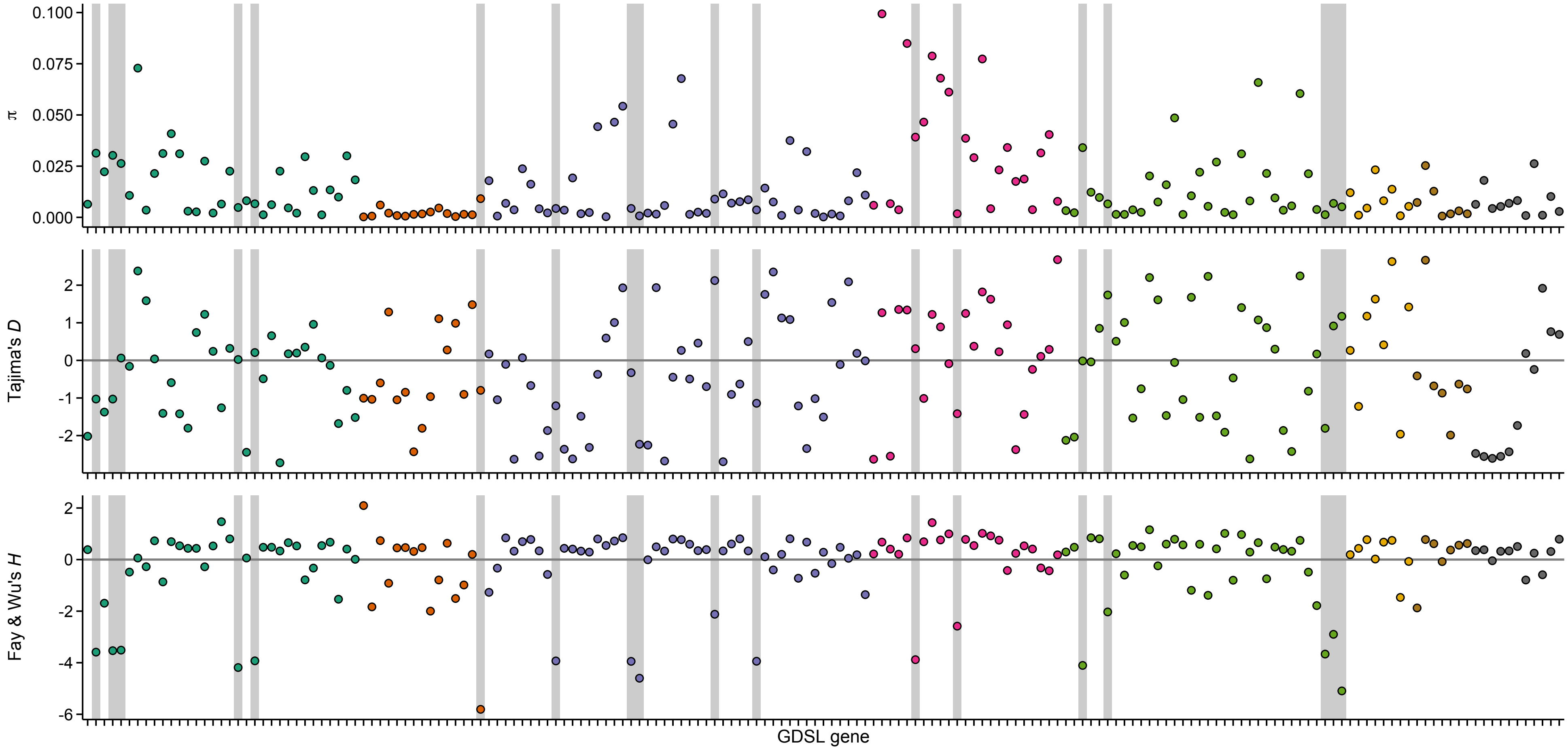


**Figure S4.** Nucleotide diversity (π), Tajima's *D*, and Fay & Wu's *H* estimated for the coding sequence of GDSL genes. Grey highlights show genes with Fay & Wu's $H < -2$. Colours correspond to clades shown in Fig. 1B of the main text.

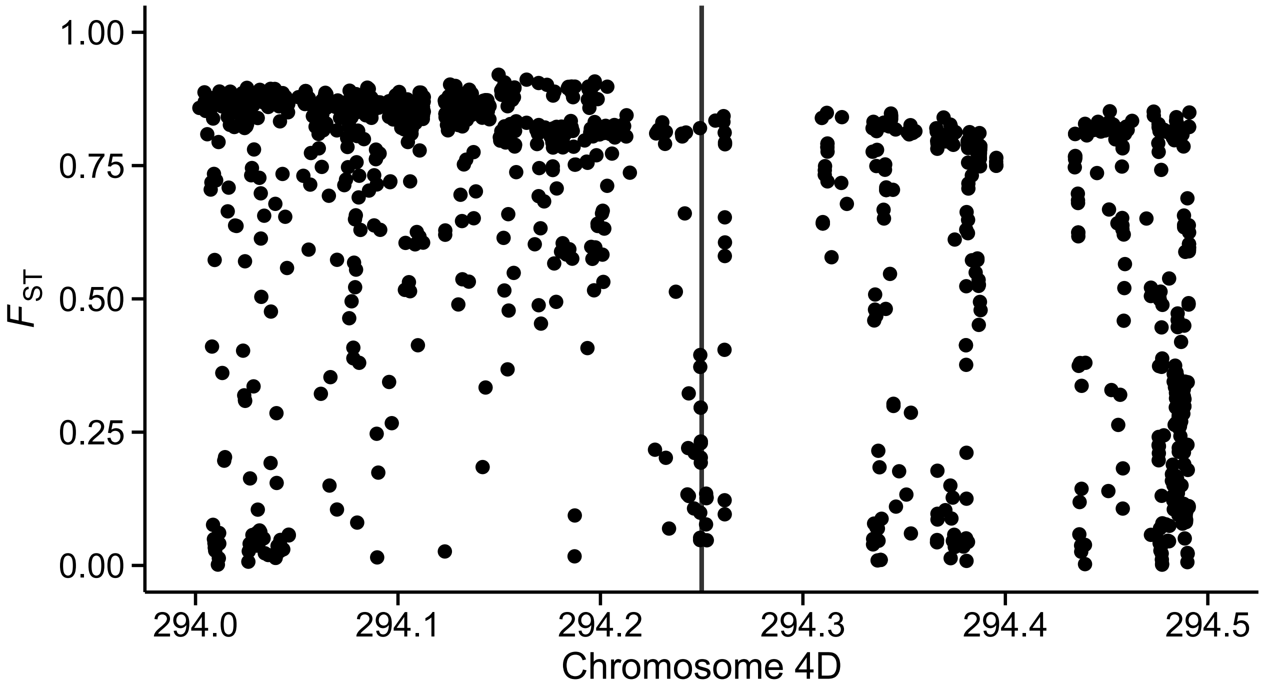


**Figure S5.** Relative nucleotide differentiation ($F_{ST}$) between *A. sativa* and *A. sterilis* in 500 kb region around gene 4DG03266120 (vertical line).
